# Sediment-associated processes account for most of the spatial variation in stream ecosystem respiration in the Yakima River basin

**DOI:** 10.1101/2024.03.22.586339

**Authors:** Vanessa A. Garayburu-Caruso, Matthew Kaufman, Brieanne Forbes, Robert O. Hall, Maggi Laan, Xingyuan Chen, Xinming Lin, Stephanie Fulton, Lupita Renteria, Yilin Fang, Kyongho Son, James C. Stegen

**Affiliations:** Pacific Northwest National Laboratory, Richland, WA 99354; Worcester State University, Worcester, MA 01602; Flathead Lake Biological Station, University of Montana, Polson, MT 59860; Washington State University, Pullman, WA 99163

## Abstract

Hyporheic zones (HZ) can contribute substantially to total stream ecosystem respiration (ER_tot_). HZ-focused process-based models may, therefore, effectively predict ER_tot_ across sites, yet this remains untested under variable environmental conditions. Here we evaluate whether spatial variation in HZ respiration predicted via a process-based model explains spatial variation in field-estimates of ER_tot_ across 33 sites in the Yakima River basin in Washington State, USA. We found that HZ respiration predictions did not explain spatial variation in field estimates of ER_tot_. To investigate further, we partitioned ER_tot_ contributions into water column respiration (ER_wc_) and sediment-associated respiration (ER_sed_). ER_sed_ contributed >50% of ER_tot_ at 88% of sites, though relative contributions varied substantially. Despite this dominance, modeled HZ respiration explained neither spatial variation in ER_tot_ nor in ER_sed_, suggesting that the HZ model alone does not capture the drivers of sediment-associated respiration across these sites. Instead, ER_sed_ spatial variation was primarily explained by gross primary production, stream slope, velocity, and total dissolved nitrogen rather than median grain size, a primary control of HZ respiration predicted by the process-based model. Consistent with recent studies, our results indicate that improving basin-scale ER_tot_ predictions requires integrating hydrologic and biogeochemical processes across hyporheic, benthic, and water column zones.

## Introduction

River corridors contribute substantial amounts of carbon dioxide (CO_2_) to the atmosphere through ecosystem respiration, a key component of aquatic ecosystem metabolism^1^. Ecosystem metabolism and carbon (C) cycling have been widely studied via two key processes: C fixation (gross primary production, GPP) and organic C mineralization (ecosystem respiration, ER)^2^.

These processes are influenced by several biotic and abiotic factors such as light, flow velocity, temperature, nutrients, microbial biomass, and hydrogeomorphology^3–6^. For example, a recent study across 222 rivers in the United States found that light availability and stream flow conditions explained variation in both annual GPP and ER^7^. Similarly, global assessments indicated that site characteristics such as land use and vegetation were more influential than latitudinal temperature effects on river metabolism^8^. These insights are derived from field estimates obtained via diel dissolved oxygen (DO) measurements. These estimates account for aerobic organisms (i.e., autotrophs and heterotrophs) and habitats (i.e., benthic, planktonic, and hyporheic zones), which together contribute to C cycling dynamics in streams and rivers at a reach scale^9^.

Beyond these field-based insights, advances in process-based modeling provide valuable understanding into stream metabolism and biogeochemical processes across scales and river corridor components (e.g., water column, benthic zone, and hyporheic zone). Recent network-scale modeling approaches have focused on representing metabolism across entire stream networks. For example, Segatto et al.^10^ developed a spatially distributed framework that couples temperature and light regimes with reach-scale ecosystem models to simulate metabolism across entire stream networks. Their approach integrates network structure, catchment vegetation, and hydrological regimes to predict organic C dynamics and ecosystem metabolism, representing streams as meta-ecosystems connected by flows of energy and materials. Such detailed process-based models have primarily been applied to relatively small catchments (e.g., 256 km² in Segatto et al.^10^) where environmental conditions and land cover are often treated as relatively homogeneous. These models also require extensive parameterization that limits their application across large, environmentally heterogeneous basins. For larger-scale efforts, modeling approaches have focused on specific river corridor components, particularly hyporheic zones (HZ, areas where groundwater and surface water mix). Models like Networks with Exchange and Subsurface Storage (NEXSS)^11^ and the River Corridor Model (RCM)^12–14^ have been used to predict variation in HZ respiration across scales ranging from individual watersheds to large basins encompassing multiple watersheds with heterogeneous biophysical conditions. These and other watershed/basin scale models rely on physical stream characteristics (e.g., sediment texture, geomorphology, discharge) to estimate hydrologic exchange fluxes that, when paired to reaction network models, provide estimates of HZ respiration rates^11–14^.

The focus on HZ modeling has been motivated by the substantial contribution of HZ processes to total stream ecosystem respiration (ER_tot_), which can range from 3-96% across different systems^15–20^. This variability suggests that HZ respiration may be a primary driver of ER_tot_ in some environments but play a secondary role in others. To understand its importance, field experiments have estimated HZ respiration via extrapolations of sediment microcosms^21,22^, as the difference between ecosystem and benthic respiration^15^, via end-member mixing analysis using in-situ solute concentration^16^ or using reactive tracers^23,24^. However, quantifying HZ respiration in the field across scales is difficult due to high spatial and temporal heterogeneity in hydrologic conditions and stream geomorphology. While process-based HZ models can overcome these field-based challenges, we lack comprehensive empirical assessments of model performance across the full range of spatial and environmental variability where HZ contributions may differ significantly.

To address this gap, we employed a hypothesis-based model-experiment (ModEx) approach to test the hypothesis that spatial variation in HZ respiration predicted via a process-based model (i.e., the RCM) explains spatial variation of field-estimates of ER_tot_. The RCM emphasizes hydrologic exchange through the HZ, including deep flow paths, while field estimates of ER_tot_ more broadly capture all surface and subsurface processes that influence DO dynamics in the water column. Our hypothesis therefore assumes that HZ respiration is the dominant driver of ER_tot_ via aerobic respiration. If that is not true across all locations or if the RCM does not represent all key processes, we may expect to reject our hypothesis. We take the perspective that valuable insights into model limitations (e.g., missing processes) are gained regardless of whether we find support for or reject the hypothesis. This hypothesis-focused approach to ModEx goes beyond data-model integration to enable iterative improvements to models and fundamental understanding.

We tested the hypothesis across 33 sites in the Yakima River basin (YRB) in Washington State, USA. Upon rejecting the hypothesis, we examined the relative contributions of water column respiration (ER_wc_) and sediment-associated respiration (ER_sed_) to ER_tot_. These additional measurements were used to test the hypothesis that ER_tot_ is driven primarily by contributions from ER_sed_, and less so by contributions from ER_wc_. Here, ER_sed_ includes HZ respiration and all other sediment-associated respiration (e.g., from the benthic zone). We found substantial variation in the fractional contribution of ER_sed_ to ER_tot_ and, in turn, investigated factors explaining variation in ER_sed_ across the basin. Our study highlights the importance of sediment-associated processes for understanding basin scale variability of ER_tot_ and emphasizes the value of process-based forward-simulation capabilities that incorporate multiple components of the river corridor, including water column processes, sediment-associated autotrophic and heterotrophic metabolism in benthic zones and HZs, and hydrologic exchange.

## Results and Discussion

Our study provided three primary outcomes: (*i*) Variation in model-predicted HZ respiration (as viewed through the river corridor model, RCM) did not explain variation in field-estimates of stream ecosystem respiration (ER_tot_) at the basin scale. (*ii*) Contributions from water column respiration (ER_wc_) and sediment-associated respiration (ER_sed_) were highly variable and ER_sed_ was the primary contributor to ER_tot_ across most sites. (*iii*) variation in ER_sed_ was primarily explained by gross primary production (GPP) and parameters reflecting the physical environment. These results suggest that there are processes driving variation in ER_tot_ at a basin scale beyond those present in the RCM. We also infer there are processes in the RCM that may not be captured in field estimates of ER_tot_. Our results provide guidance for which processes could be included in basin-scale process-based models, such as the RCM, to support research efforts focused on understanding patterns and dynamics of ER_tot_ and ER_sed_ across large environmental gradients.

*Spatial variation in RCM-predicted HZ respiration did not explain spatial variation in field-estimates of ER_tot_ across the YRB*.

Estimates of time-averaged ER_tot_ for the deployment period across sites (n = 45, Methods – Sensor deployment and data collection) were highly variable and ranged from −21.50 to +9.26 g O_2_ m⁻^2^ day⁻^1^ (Fig. 1a). After accounting for biologically impossible values (see below), the range of biologically relevant time-averaged ER_tot_ (n = 33, Fig. S1a) values used for the remainder of analyses ranged from −21.50 to −0.12 g O_2_ m^⁻2^ day^⁻1^. We could not estimate metabolism (i.e., GPP and ER_tot_) in 3 sites that failed to meet the assumptions of one station open channel O_2_ budgets due to slower flows and lack of turbulence^25^ (Fig. S2a). In addition, out of the 45 sites where we were able to estimate metabolism, 12 sites displayed positive estimates of ER_tot_ (Fig. S2a). These had dissolved O_2_ (DO) that rarely went under 100% saturation, which indicates high gas exchange rates potentially due to too much turbulence and bubbles at higher internal pressure relative to the atmosphere^26^. Oversaturation hinders the estimation of ER_tot_ using open channel DO time series because we have no simple way of knowing what the concentration of DO would be in the river in the absence of any biological activity^27^. We removed sites with positive ER_tot_ values from our analysis because they have no biological meaning, and their magnitude reflects misspecification due to bubble-mediated gas exchange and not a biological process.

**Figure 1.**
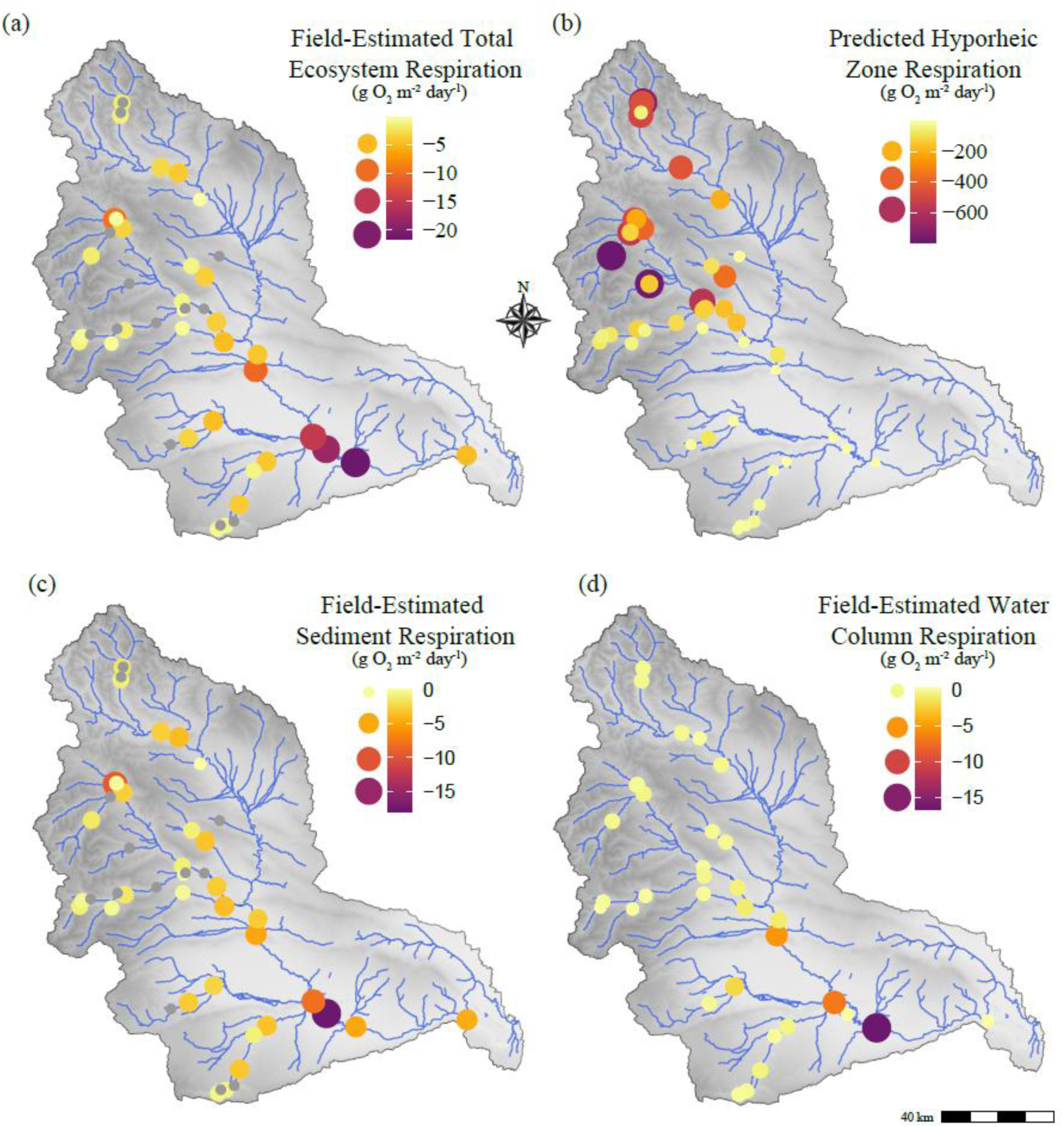
Map displaying Yakima River basin sites investigated in this study and associated respiration rates. (a) Field-estimates (n = 48) of total stream ecosystem respiration rates (ER_tot_). (b) River Corridor Model (RCM, n = 46) predicted hyporheic zone (HZ) respiration rates^14^. (c) Field-estimates (n = 48) of sediment-associated respiration rates (ER_sed_). (d) Field-estimates (n = 48) of water column respiration rates (ER_wc_). Color and symbol size are used to indicate magnitude of respiration rate. Grey dots in panels a and c indicate sites where ERtot or ER_sed_ could not be estimated or where ER_tot_ was positive.

Field-estimates of ER_tot_ diverged from RCM-predicted HZ respiration, both quantitatively and qualitatively. The fastest ER_tot_ rates occurred in the lower region of the YRB (Fig. 1a), in locations with shallow topographic gradients and high order streams (i.e., topographic slopes of 0.0004 - 0.002 and stream orders of 5-7). The fastest HZ respiration rates predicted by the RCM were, in contrast, in upper parts of the YRB (Fig. 1b). RCM-predicted HZ respiration values (n = 46) ranged from −800 to −0.024 g O_2_ m⁻^2^ day⁻^1^. The RCM-predicted rates were much faster than estimates of ER_tot_ highlighting fundamental differences in the processes represented in the RCM vs. processes that influence ER_tot_ field estimates.

Our ModEx approach focuses on evaluating whether current process-based representations of HZ respiration, as implemented by the RCM, can explain observed spatial variation in field estimates of ER_tot_. While improving the RCM or developing alternative models is beyond the scope of the current study, our results provide valuable guidance for future model development by identifying which processes may be missing or inadequately represented in models trying to understand basin-scale ER_tot._ Therefore, we use RCM-predicted values as the best current representation for spatial variation in HZ respiration rates across the YRB. This approach allows for evaluating a qualitative relationship between the spatial variation of RCM-predicted HZ respiration and field-based ER_tot_.

Spatial variation in RCM-predicted HZ respiration had a weak negative relationship with spatial variation in ER_tot_. Before doing associated analyses and given the quantitative difference between RCM predictions and ER_tot_, we standardized rates within each dataset as z-scores, whereby each dataset had a mean of zero and standard deviation of one. This does not change quantitative relationships within each dataset or the shape of the data distribution; it only puts the two datasets into the same quantitative scale. Following this normalization, we regressed RCM-predicted HZ respiration z-scores against z-scores of field-estimates of ER_tot_ (Fig. 2).

**Figure 2.**
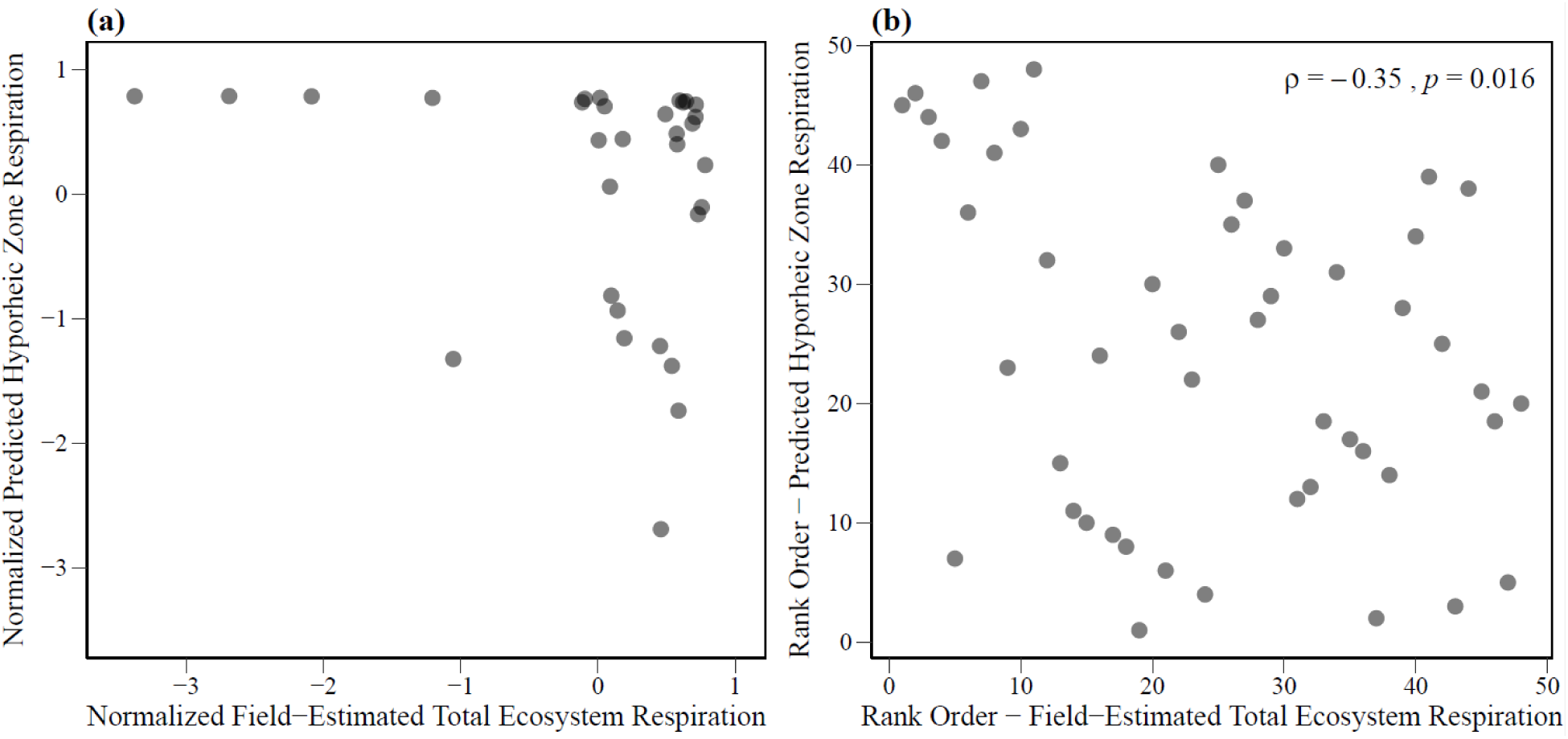
Normalized model-predicted hyporheic zone (HZ) respiration rates do not increase with field-estimates of total stream ecosystem respiration rates (ERtot; n = 31). (a) Z-score normalized data suggests no relationship. Data distributions are skewed (i.e., not normal) and thus violate assumptions of parametric regression, so no statistics are given. (b) Spearman rank-order correlation reveals a weak, negative relationship. The Spearman correlation statistics are provided on the panel.

These data violated the normality assumption for conducting a parametric regression, yet no relationship is apparent in the data (Fig. 2a). While Spearman rank-order correlation appeared to show a negative relationship, it was weak enough to be considered likely irrelevant (Fig. 2b). Rank-order correlation was used because the data violated normality assumptions required for parametric analysis, and plotting the ranked data makes the monotonic relationship between the variables more apparent than the normalized values. The negative direction and weakness of the relationship indicate, however, that HZ respiration—as represented in the RCM—is not a primary factor explaining spatial variation of ER_tot_ in the YRB. The lack of a strong positive relationship could be for several reasons, such as ER_tot_ not including respiration occurring across the full HZ spatial domain that is represented in the RCM, and the RCM not including all the river corridor components and drivers influencing ER_tot_ estimates.

Field-estimates of ER_tot_ are based on DO dynamics that reflect the reach-scale integrated outcome of processes occurring across all system components that are hydrologically connected to the active channel^9^. This estimate includes respiration associated with subsurface sediments along relatively short HZ flow paths. These HZ flow paths must be fully contained within the upstream reach that influences observed DO dynamics at a given monitoring location. If the HZ flow paths are longer, they should not influence observed DO and thus do not influence estimated ER_tot_. The length of the reach that a given DO sensor integrates varies across streams and has been estimated to be roughly three times the turnover length of O ^9^.

The RCM emphasizes hydrologic exchange through the HZ, including deep flow paths, thus it is plausible that the RCM captures respiration along flow paths that are longer and deeper into the HZ than what the DO sensors integrate across. If so, respiration in long flow paths—that are captured in the RCM—could lead to the lack of positive correlation between RCM-predicted HZ respiration and field-estimates of ER_tot_. We suspect, however, that this might not be the primary reason for the lack of a relationship because both RCM-predicted HZ respiration and field-estimates of ER_tot_ account for short HZ flow paths. These shorter flow paths are likely to be within the most biogeochemically active sediments, which should in turn be the primary drivers of total HZ respiration^28^. We postulate that the lack of a relationship (Fig. 2) and divergence in spatial variation (Fig. 1a, b) is related to differences in which biogeochemical processes are captured in the two methods.

The RCM represents hydrologic exchange through the HZ and sediment-associated respiration that is fueled by reactants delivered by that exchange^14^. In the model, reactants delivered to HZ sediments are not influenced by primary producers (e.g., photoautotrophic algal biofilms) and the respiration contributed by primary producers is also not accounted for. Respiration occurring by heterotrophs and photoautotrophic primary producers in the water column is also not part of the RCM. All these processes can influence observed DO dynamics^29–31^ and are thus implicitly accounted for in field-estimates of ER_tot_. This highlights that field estimated ER_tot_ is influenced by several processes that are not represented in the RCM. This is not a critique or rejection of the RCM, rather, it is a recognition that the divergence in patterns between RCM-predicted HZ respiration and field-estimates of ER_tot_ are likely due, at least in part, to a difference in the processes captured by each method.

In addition to the differences in captured processes, RCM-predicted HZ respiration and ER_tot_ may not be correlated if processes in the RCM do not strongly influence ER_tot_, if data inputs to the RCM misrepresent the system, or if there are technical issues associated with estimating ER_tot_. The ER_tot_ estimates used in our analysis only included sites where the assumptions of one station open channel O_2_ budgets were met and where streamMetabolizer results converged (n = 33). Moreover, since our estimates of ER_tot_ largely overlapped with literature values^7,32^ (Fig. 3a), it is, likely that our ER_tot_ estimates are technically robust. We infer that technical issues with our ER_tot_ estimates are not a cause for the lack of correlation with RCM predictions. Thus, the lack of correspondence in patterns suggests that to use the RCM, or other processed-based models, in future studies to help understand ER_tot_ it may be useful to expand the breadth of represented processes (e.g., as in Segatto et al.^10^). Available field estimates of ER_tot_ allow us to guide which processes could be most important to include in the RCM and, more generally, to investigate factors associated with spatial patterns in ER_tot_. To these ends, we next examine the relative influence of water column- and sediment-associated processes over ER_tot_, which provides conceptual understanding of system function and evaluates the need for each of these in an expanded version of the RCM.

**Figure 3.**
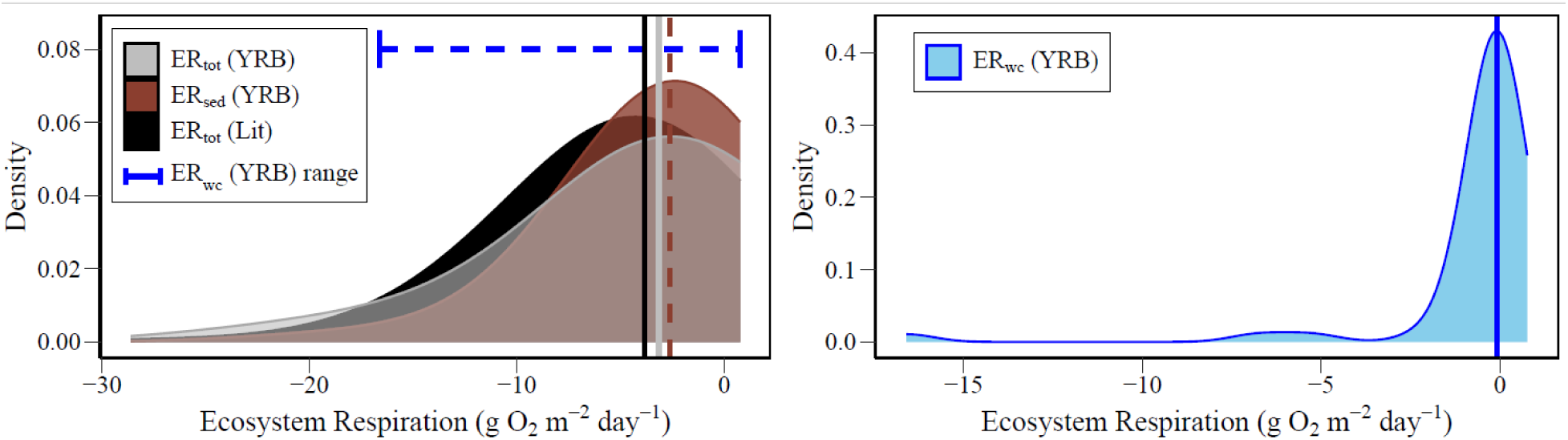
Density plots comparing respiration rates for different river corridor components within the Yakima River basin (YRB) as well as literature values. (a) Density plot for YRB total ecosystem respiration from this study (ER_tot_, grey, n = 33) and from rivers and streams across the Contiguous United States (CONUS) (n = 222 estimated by Appling et al.^32^ and Bernhardt et al.*^7^* (ER_tot_(Lit), black), as well as YRB ER_sed_ (brown, n = 33) and the range of YRB water column respiration rates (ER_wc_, blue n = 48); (b) Density plot for YRB ER_wc_ (blue). The vertical lines are the medians for each distribution.

### Relative contributions of ER_sed_ and ER_wc_ to ER_tot_ varied substantially across sites

Our ER_wc_ data, estimated via *in-situ* dark bottle incubations (n = 48), varied greatly across the basin and ranged from −16.60 to +0.772 g O_2_ m^-2^ day^-1^ (Fig. 3b). Contrary to the data cleaning decisions made for ER_tot_, we chose to interpret positive ER_wc_ because they reflect instrument noise paired with extremely slow respiration. In this situation the rate at which DO was consumed was not fast enough to overcome instrument error and resulted in a small positive ER_wc_ measurement. If these values were excluded it would bias the dataset away from locations with very slow ER_wc_ rates. Across all 33 field locations with ER_tot_ estimates, ER_wc_ varied greatly and had fractional contributions of ER_wc_ to ER_tot_ spanning 0-100% (Table 1, Fig. S3a). The range of measured ER_wc_ values spanned a substantial portion of ER_tot_ literature values (Fig. 3a). These results highlight the variable environments in the YRB and suggest that ER_wc_ can be of similar magnitude to ER_tot_ across rivers in the contiguous United States (CONUS).

**Table 1.** Summary statistics (mean, median, and range) for ER_tot_ (g O_2_ m^-2^ day^-1^), ER_wc_ (g O_2_ m^-2^ day^-1^), ER_sed_, (g O_2_ m^-2^ day^-1^), and ER_wc_ and ER_sed_ contributions to ER_tot_ (percent).

|  | Mean | Median | Range |
| --- | --- | --- | --- |
| $ER_{tot}$ | -4.14 | -3.15 | (-21.50, -0.12) |
| $ER_{wc}$ | -0.725 | -0.0845 | (-16.60, 0.772) |
| $ER_{sed}$ | -3.05 | -2.62 | (-17.96, 0) |
| $ER_{wc}$ (%) | 8.4% | 17.7% | (0%, 100%) |
| $ER_{sed}$ (%) | 82.3% | 91.7% | (0%, 100%) |

As discussed in Laan et al.^33^, ER_wc_ in the YRB spans values that are similar to other streams and rivers around the world^34–36^. Variable ER_wc_ rates in the YRB could be due to variable amounts of total suspended solids (TSS), nutrient concentrations, and changes in depth and surface water residence time, moving from headwaters to larger order streams^33,34,37^. Higher TSS increases surface area for microbial attachment which can lead to higher metabolic rates within the planktonic compartment of the river^38–40^. Higher nutrient concentrations, particularly nitrogen and phosphorus, support increased microbial biomass and metabolic activity in the water column by alleviating nutrient limitation^41,42^. Additionally, increasing depth and longer surface water residence times in larger streams create conditions that favor water column processes over benthic processes due to larger water volume-to-benthic area ratios and more complete processing of organic matter by planktonic communities^36,43,44^.

Moreover, our results indicate that respiration contributions from ER_wc_ can vary widely across the YRB during base-flow conditions. The median ER_wc_ contribution was 17.7%, though ER_wc_ contributed >50% to ER_tot_ at 4 sites (12% of all sites), with contributions ranging from 62-100% at these locations. These sites corresponded to 4^th^ and 5^th^ order streams with the slowest ER_tot_ rates (−0.12 to −1.45 g O_2_ m⁻² day⁻¹) across our dataset. While ER_wc_ and ER_tot_ rates at these locations were slow, the water column was responsible for nearly all of the observed respiration. These findings align with previous studies finding large ER_wc_ contributions. For example, working in mid-sized rivers, Reisinger et al.^36^ found that ER_wc_ contributed an average of 35% to ER_tot_, and up to 81% in rivers with slow ER_tot_. Similarly, the rates of other water column processes, like nutrient uptake and denitrification, may increase with increasing river size, as material processing shifts from being benthic-dominated to water-column dominated in rivers from 5^th^ to 9^th^ order^42,45–47^. These patterns suggest that the relative importance of ER_wc_ is governed by several stream characteristics, some of which co-vary across the basin.

We further examined respiration contributions from river corridor components beyond the water column (i.e., benthic/streambed sediments and HZs that are hydrologically connected to the active channel). We define the respiration that comes from these integrated components as sediment-associated respiration (ER_sed_). ER_sed_ was calculated as the difference between ER_tot_ and ER_wc_ at each site with negative ER_tot_ (n = 33) assuming zero ER_wc_ rates if ER_wc_ values were positive (n = 12). This approach allows for the full range of variability in respiration contributions, from sites where ER_wc_ dominates (i.e., ER_sed_ contributions approach zero) to sites where ER_sed_ dominates (i.e., ER_wc_ contributions approach zero). ER_sed_ values ranged from −17.96 to 0 g O_2_ m^-2^ day^-1^ (Table 1). The relative contribution of ER_sed_ to ER_tot_ spanned from 0-100%, with 29 out of 33 locations (88% of locations) showing ER_sed_ contributions exceeding 50% of ER_tot_ (Table 1, Fig. 4a). The strong contribution of ER_sed_ to ER_tot_ across the YRB aligns with previous studies that have found large contributions of HZ and benthic sediments to ecosystem respiration^16,48–50^. It is important to note that our ER_sed_ measurements integrate multiple ecosystem components, including the HZ, benthic biofilms, and submerged macrophytes, whereas ER_wc_ specifically represents respiration from organisms suspended in the water column.

**Figure 4.**
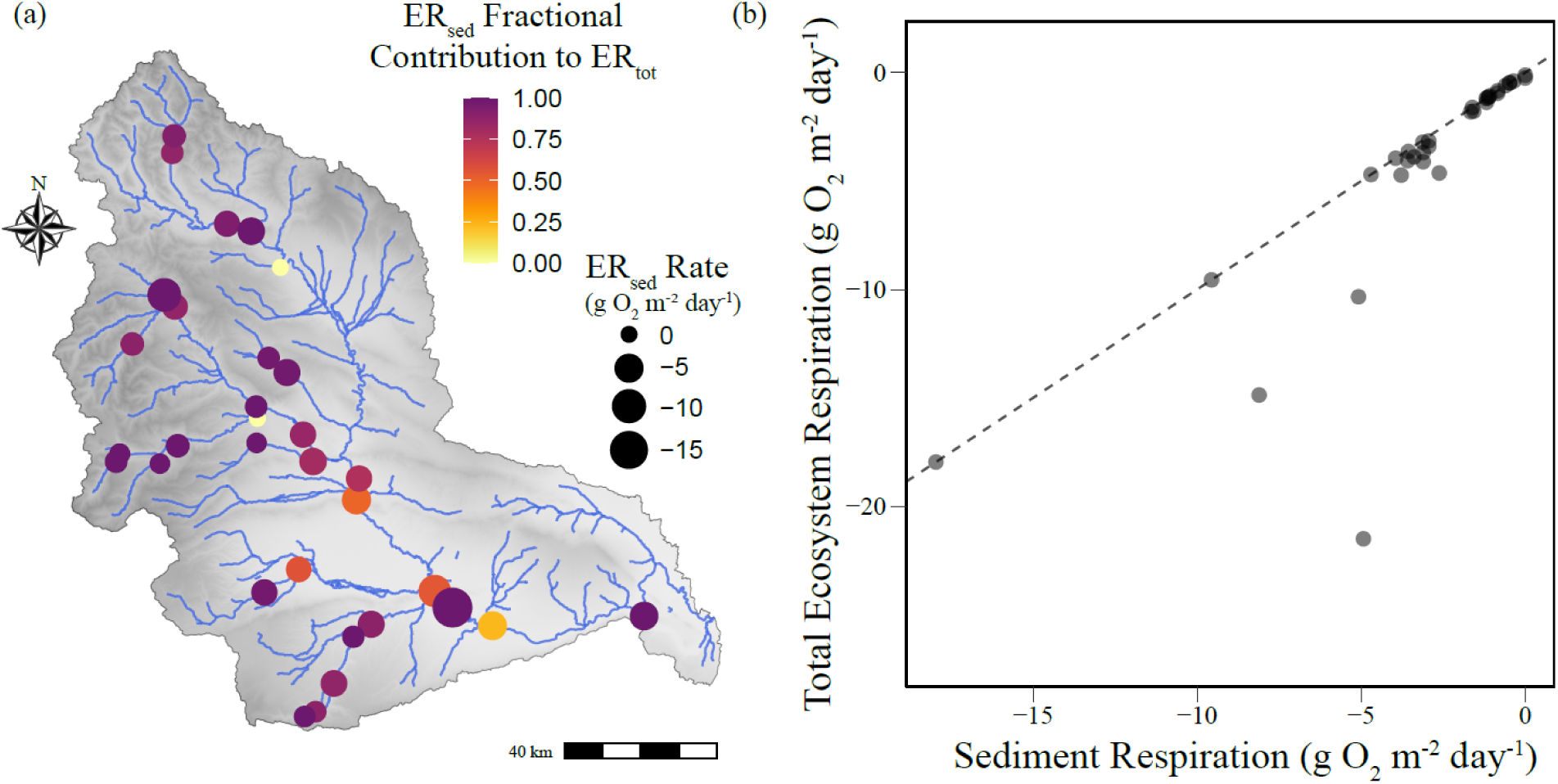
ER_sed_ contributions to and correspondence with ER_tot_. (a) Map displaying the fractional contributions of sediment-associated respiration (ER_sed_) to total ecosystem respiration (ER_tot_) across the YRB (n = 33). Color is used to indicate the magnitude of the fractional contribution while symbol size is used to indicate magnitude of ER_sed_ respiration rate. (b) Scatterplot to display the contributions of ER_sed_ to ER_tot_. The grey dashed line is the one-to-one line.

While ER_sed_ contributions accounted for most of ER_tot_ across the YRB, no correlation was observed between ER_sed_ and RCM-predicted HZ respiration (Fig. S4). ER_sed_ was fastest in lower parts of the YRB, while the RCM predicted the fastest HZ respiration to be in upper parts of the YRB (Fig.1b, c). The correspondence between ER_sed_ and ER_tot_ reflected a strong contribution of ER_sed_ to ER_tot_, with some notable exceptions at sites with substantial ER_wc_ contributions (Fig. 4b). This finding is consistent with previous studies. For example, Naegeli and Uehlinger^15^ found that ER_sed_ contributed 76–96% of ER_tot_ in a gravel bedded river using closed chamber measurements. Similarly, Fellows et al.^22^ reported large HZ contributions to ER_tot_ (40-93%) in two headwater streams under baseflow conditions. In contrast, Battin et al.^16^, found lower HZ contributions (41% of ER_tot_) in a third-order piedmont stream, while Roley et al.^43^ observed that water column processes dominated ER_tot_ in the mainstem Columbia River (9^th^ order river).

These varying results, together with our findings, demonstrate that ER_sed_ contributions can span a wide range across river networks. While these studies provide valuable insights into specific stream reaches, they highlight a critical knowledge gap in our understanding of what drives ER_sed_ variability at the basin scale. Our dataset from the YRB provides an opportunity to investigate potential drivers of spatial variation in ER_sed_ across a river network with heterogeneous environmental conditions.

### Spatial variability in ER_sed_ was best explained by primary production and physical settings

To investigate factors explaining spatial variation in ER_sed_, we identified variables known from previous work to influence ER_tot_ (e.g., gross primary production; GPP) and variables that influence HZ respiration (e.g., median grain size)^7,8,14^. We did not include RCM-predicted HZ respiration as an explanatory variable because it was strongly correlated with median grain size (i.e., d50), and because we did not find a positive correlation between RCM-predicted HZ respiration and ER_tot_ or ER_sed_ (Fig. 2, Fig. S2). Our analysis showed that GPP, stream slope, stream velocity, and total dissolved nitrogen (TDN) explained the most variation in ER_sed_ (Table 2). Other potential drivers (i.e., land cover classes, median grain size, dissolved organic carbon, depth and water temperature) had smaller regression coefficients (< 0.1) and were interpreted as not being as important. The directionality of these relationships further showed that the fastest ER_sed_ rates had the fastest GPP rates (Table 2). This finding is generally consistent with the observation from Bernhardt et al.^7^ who found that ER_tot_ was most strongly linked to light availability, which should influence GPP. Additionally, faster ER_sed_ rates were observed in reaches with shallower topographic gradients (full slope range: 0.0004-0.027), lower velocity (full velocity range: 0.29-1.44 m s^-1^), and higher TDN concentrations (full TDN range 0.05-1.47 mg N L^-1^). Flow velocity likely influences ER_sed_ due to its control over biofilm scouring, physical structure, and functional dynamics^51,52^. Moreover, stream slope influences flow velocities, sediment transport, and nitrogen concentrations, which can all impact biofilm community structure and C uptake ^53,54^. Our results suggest that together these variables can have a substantial effect on benthic metabolism and, in turn, ER_sed_.

**Table 2.**
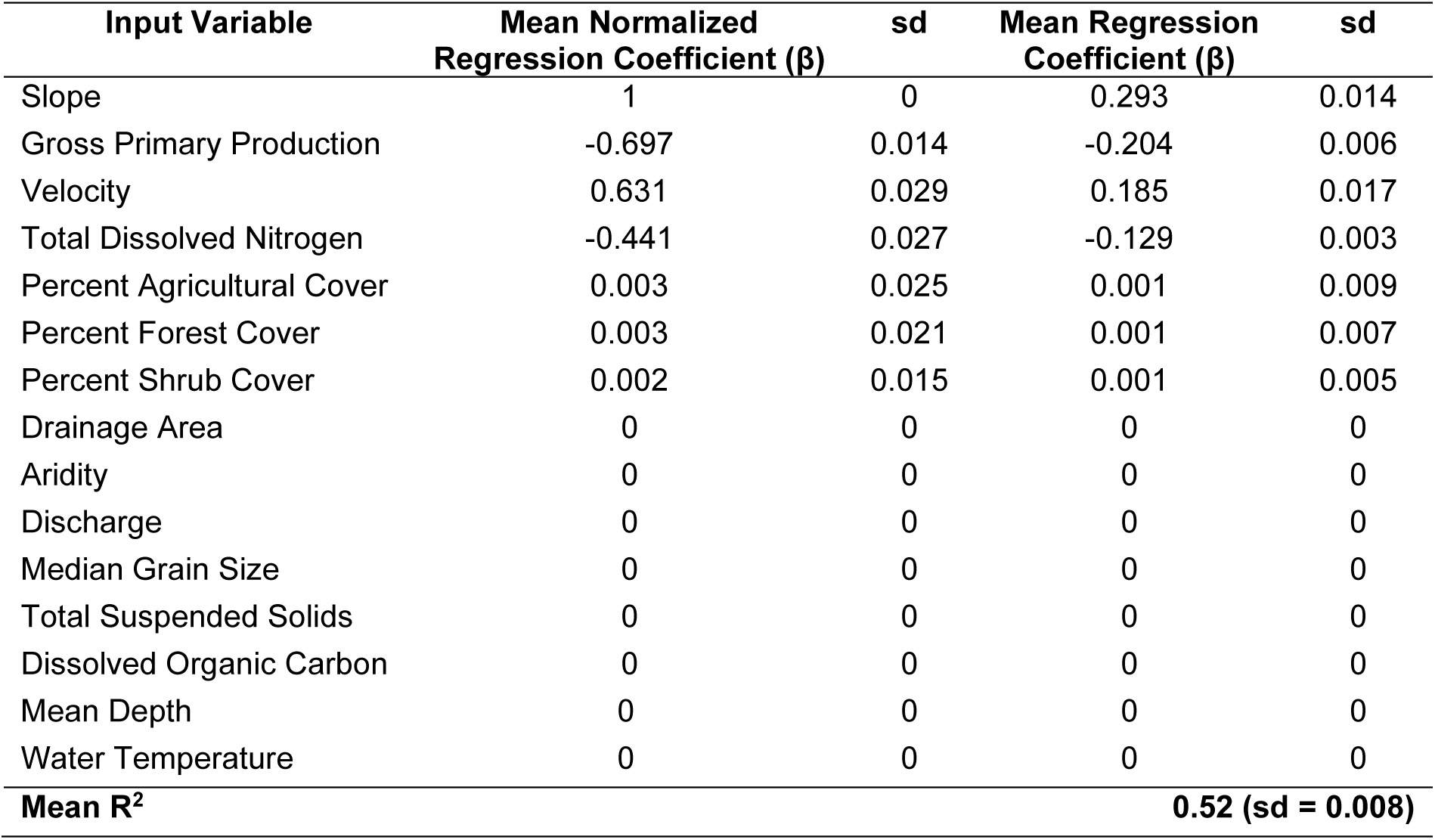
Regression coefficients for LASSO models explaining variation in ER_sed_. All input variables from 33 sites were cube root transformed to reduce the impact of high leverage points in the regression analysis and standardized as z-scores to enable direct comparison of the regression coefficients. The LASSO regressions were performed over 100 seeds. Reported mean regression coefficients (β) and R^2^ were averaged across the 100 seeds. Normalized regression coefficients were calculated by dividing each β coefficient by the maximum β coefficient in each seed. Standard deviation (sd) was calculated for each coefficient over the 100 seeds. A value of 0 indicates that the variable was included as a candidate predictor but was not selected by the model (i.e., its coefficient was shrunk to zero by LASSO regularization).

In addition, ER_sed_ and ER_tot_ covaried linearly with GPP (Fig. 5) providing insight into net ecosystem production and the metabolic balance of the stream. This information can be discerned by examining the position of sites relative to the one-to-one line in Fig. 5. Points below the dashed line indicate more respiration than primary production (i.e., net heterotrophy; ER_tot_ or ER_sed_ > GPP) and points above the line indicate more primary production than respiration (i.e., net autotrophy; GPP > ER_tot_ or ER_sed_). Our results showed that 22 sites were net autotrophic and 11 sites were net heterotrophic. This finding contrasts the general paradigm that stream ecosystems are predominantly heterotrophic, where respiration is sustained by allochthonous C inputs, contributing to rivers being net sources of CO_2_ to the atmosphere^7,9,55–57^. In autotrophic conditions, rivers fix more carbon than they respire, potentially storing or exporting this excess carbon downstream^9^. Larger number of sites exhibiting net autotrophy in the YRB might be due to summer baseflow conditions when light availability is maximized, combined with stable flow conditions that promote primary production^58^. Similar patterns of autotrophy have been observed during baseflow in agricultural systems^59^, montane streams with some seasonal and year-to-year variability ^60^, and managed pastoral streams^61^.

**Figure 5.**
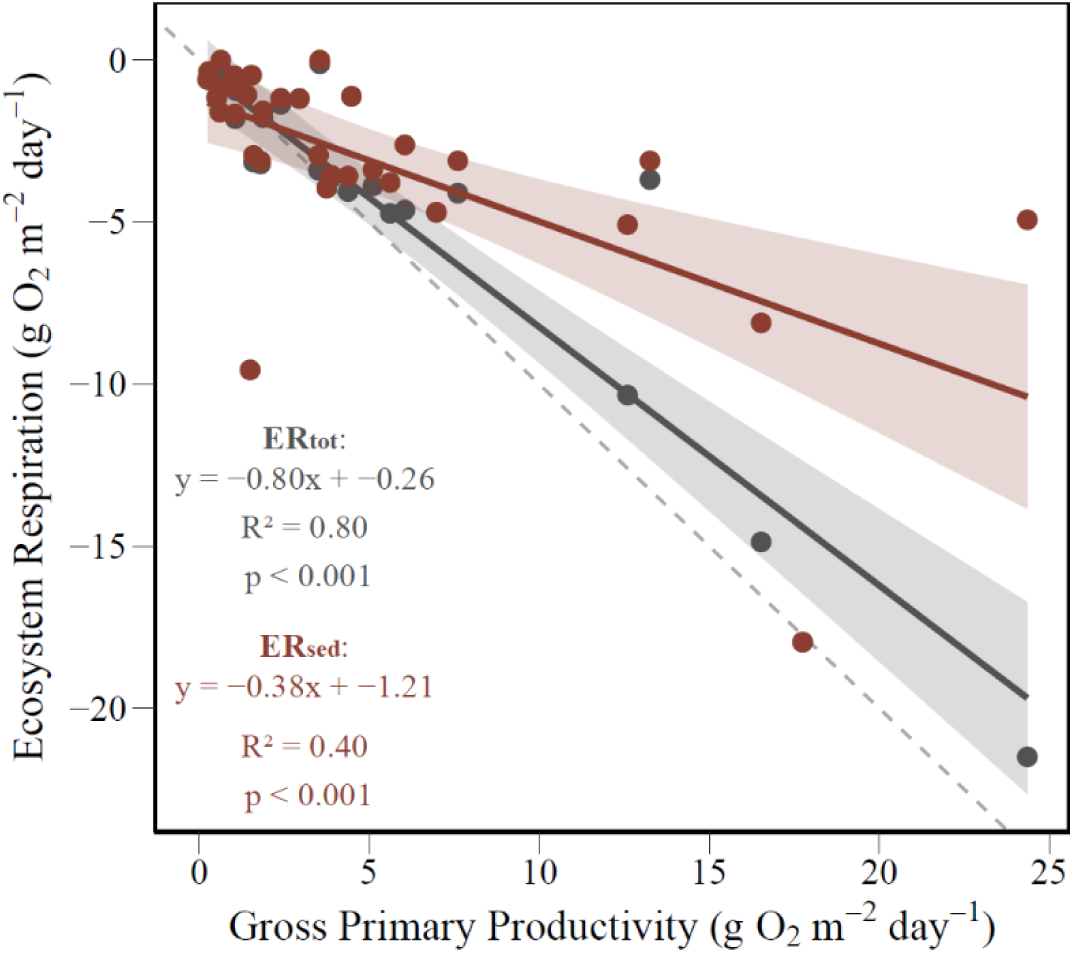
ER_sed_ and ER_tot_ as a function of GPP. Scatterplot with linear regression lines depicting the relationship between GPP (x-axis) and field estimates of sediment respiration (ER_sed_, brown, y-axis) and field estimates of total respiration (ERtot, dark grey, y-axis). The grey dashed line is the one-to-one line with slope of –1.

The relatively large LASSO coefficient for GPP (the largest coefficient after slope, Table 2), and faster ER_sed_ (more negative) with increasing GPP (Fig. 5), suggest an important link between sediment-associated primary production and ER_sed_. Primary producers in this case are likely to include both algal biofilms attached to benthic/streambed sediments and submerged macrophytes rooted in sediments. We infer that both types of primary producers likely provide important direct and indirect contributions to ER_sed_. Direct contributions include phototroph respiration, and indirect contributions include providing resources (e.g., organic exudates) to sediment-associated heterotrophs^2,62^. This link between ER_sed_ and GPP in the YRB is conceptually consistent with Bernhardt et al.^7^ who identified light as a primary explanatory variable for ER_tot_ across CONUS. Further, in the YRB, low-gradient reaches with slow moving water had faster ER_sed_ (Table 2), suggesting residence time was long enough to enable significant reactions to occur, thereby enhancing ER_sed_. These relationships suggest a complex coupling between physical, chemical, and biological drivers linked to sediment-associated primary production, residence times, and reaction rates^63–65^. Together, these results indicate that ER_sed_ hot spots^66^ or control points^67^ are more likely to occur in locations with shallow topographic gradients, times of slow flow causing long residence time, and times of fast GPP causing short reaction times. These factors are likely predictable from physical features of stream networks, thereby providing opportunities to develop mechanistic or data-driven models of ER_sed_ hot spots/control points

## Conclusions

Spatial variation of ER_sed_ largely corresponded to ER_tot_ patterns, thus understanding basin-scale ER_tot_ requires deeper understanding of controls on ER_sed_ spatial variation. Our analysis revealed that ER_sed_ across the YRB was primarily explained by slope, GPP, velocity and TDN with the fastest ER_sed_ rates occurring in reaches characterized by high GPP, gentle slopes, slower velocities, and elevated TDN concentrations. These relationships suggest a complex coupling between physical, chemical, and biological drivers linked to sediment-associated primary production, residence times, and reaction rates. Further, the lack of positive correlation between RCM-predicted HZ respiration and field-estimates of ER_sed_ and ER_tot_ highlights a need to extend conceptual and process-based models that aim to capture and predict active channel respiration to represent all processes that have major influences over those rates. The RCM relies heavily on several spatially variable inputs, including median grain size (i.e., d50) and dissolved organic carbon (DOC) concentrations. These input data layers, particularly d50, are generated from sparse field measurements and may not accurately reflect local conditions in the basin^68^. If the spatial patterns in d50 within the input data deviate from true spatial patterns in d50, that will cause errors in predicted spatial patterns of HZ respiration. Additionally, the model does not consider the impact of sediment grain size on reaction rates, only on exchange rates.

To understand patterns in ER_tot_ and ER_sed_ across sites we propose that it is not enough to focus on HZ respiration as influenced primarily by hydrologic exchange processes. Our results indicate a need to explicitly represent primary producers and heterotrophs in both the water column and associated with sediments. Current models, such as the RCM, can be modified to meet this need by including: (1) sediment-associated primary production and its coupling with heterotrophic respiration; (2) water column metabolic processes, particularly for larger rivers where ER_wc_ can dominate; and (3) better constraining data layers (e.g., d50) through targeted field measurements^69^. Recent advances in network-scale biogeochemical modeling (e.g., Segatto et al.^10^) demonstrate the value of incorporating detailed process representations across stream networks. These approaches have primarily been applied to smaller, more homogeneous catchments and require integration with three-dimensional physics-based flow and transport models to capture the hydrologic complexity and environmental heterogeneity of large basins like the YRB.

Developing integrated modeling capabilities for large-scale studies requires iterative evaluation of model performance against empirical observations across a broad range of conditions. We refer to this approach as hypothesis-based model-experiment (ModEx) iteration, and this study represents a key step in this approach. Hypothesis-based ModEx goes beyond data-model integration whereby models are used to pose new hypotheses that are tested with new data, leading to modified hypotheses that guide follow-on model development, and so on. We encourage the application of this approach in other basins and across other flow conditions to evaluate the transferability of our outcomes. This is key to building predictive models that are useful across Earth’s watershed systems.

## Methods

### 1. Site selection

We selected the Yakima River basin (YRB) in Washington State, USA as a test basin for this study. The basin drains an area of 15,941.4km^2^ and spans forested mountainous regions in the west to arid valleys and plains in the east (Fig 6). The basin has variable land covers and land uses dominated by dry rangeland, forest, and agriculture^70,71^. Annual precipitation ranges from 203-356 cm in the western parts of the basin to 25 cm in the eastern parts of the basin ^72^.

**Figure 6.**
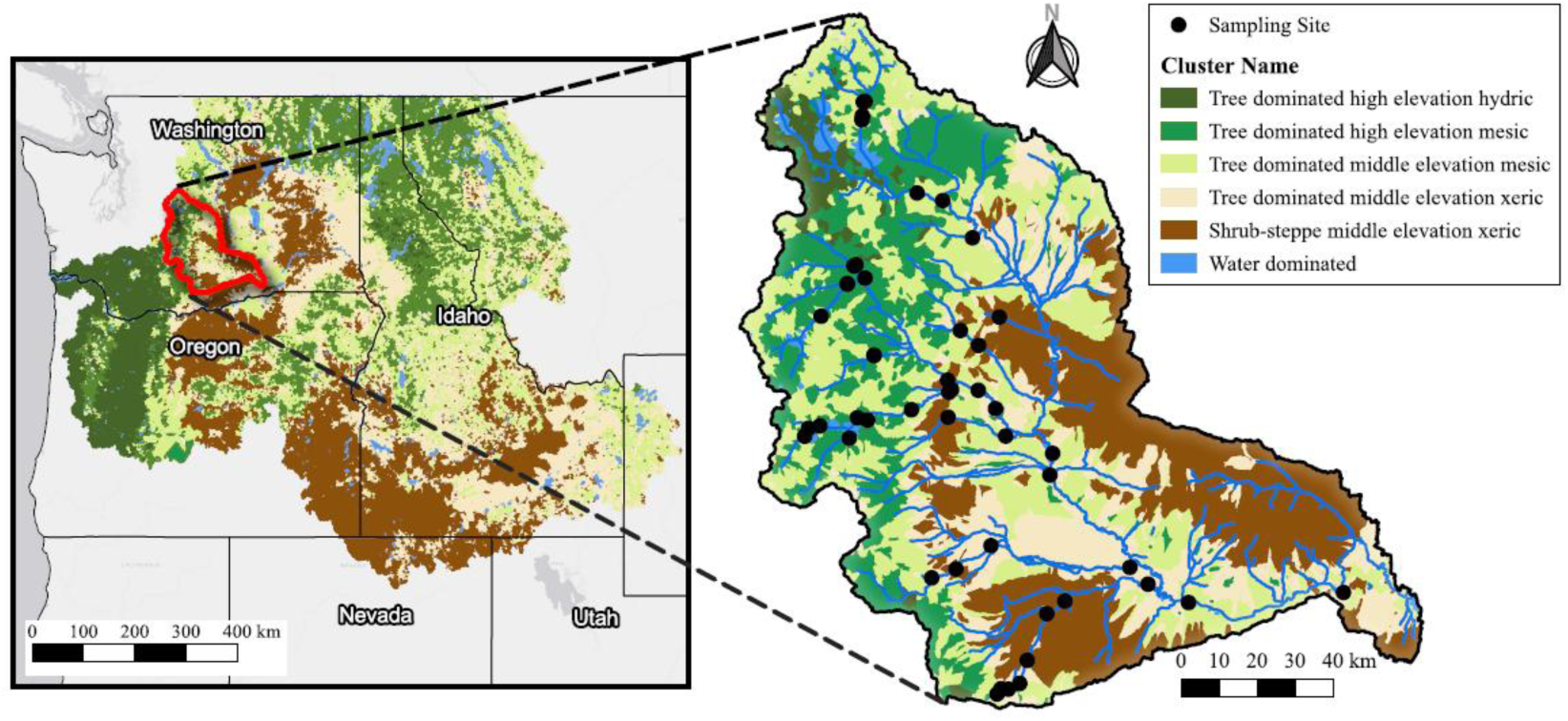
Yakima River basin sampling sites superimposed on a map of catchment cluster analysis results.

Field sites spanned a range of drainage areas, land cover types, physical settings, and previously published model-predicted HZ respiration rates^14^. Site selection was designed to span the range of biophysical conditions that influence HZ respiration as predicted by the RCM, including variations in drainage area, slope, sediment grain size, and land cover. This approach ensures that our results are not biased toward particular environmental conditions and provides a robust test of model predictions across a range of environmental settings. Because model-predictions were performed at a reach scale for the Columbia River basin, we conducted a clustering analysis following methods previously described by Laan et al.^33^ to locate regions in our test basin with representative characteristics from the Columbia River basin (Fig. 6). Cluster analysis used 15 environmental variables including elevation, precipitation, temperature, land cover percentages, and topographic characteristics following methods in Laan et al^33^. We then combined the clustering analysis results with logistical considerations regarding site access, safety, and model-predicted HZ respiration rates to select 48 sites that span a range of clusters, model-predicted respiration rates and drainage areas across the basin (Fig. 6). The selected sites are collectively representative of 96% of the drainage area across six major biophysical clusters in the larger (∼670,000 km^2^) Columbia River Basin (Table 3), supporting the transferability of our findings due to unbiased and representative sampling. Table 3 shows a summary of the cluster analysis results as well as number of sites in the YRB per cluster.

**Table 3.** Cluster analysis results across the Columbia River basin and Yakima River basin with similar biophysical and hydrologic characteristics and the number and percentage of sites in each.

| Cluster | Cluster Name | CRB Drainage Area (%) | YRB Drainage Area (%) | No. YRB Sites Per Cluster | Percent YRB Sites Per Cluster (%) |
| --- | --- | --- | --- | --- | --- |
| 1 | Tree dominated high elevation mesic | 22.7 | 27.4 | 12 | 25.0 |
| 2 | Water dominated | 3.4 | 2.2 | 0 | 0 |
| 3 | Tree dominated high elevation hydric | 7.3 | 2.4 | 2 | 4.2 |
| 4 | Shrub-steppe middle elevation xeric | 25.4 | 28.1 | 5 | 10.4 |
| 5 | Tree dominated middle elevation mesic | 17.4 | 17.3 | 15 | 31.3 |
| 6 | Tree dominated middle elevation xeric | 23.9 | 22.6 | 14 | 29.2 |

*"CRB Drainage Area" is the percentage of the total drainage area of the Columbia River basin that was classified in each cluster.*
*"YRB Drainage Area" is the percentage of the total drainage area of the Yakima River basin that was classified in each cluster.*
*"YRB Sites Per Cluster" is the total number of field sites in the Yakima River basin (n = 47) located in each cluster. "Percent YRB*
*Sites Per Cluster" is the percentage of the total number of sampling sites in the Yakima River basin located in each cluster.*

Geospatial variables for each site were extracted primarily from the Environmental Protection Agency StreamCat Dataset and NHDPlusV2.1 using a custom R script^73^. Land use/land cover data for all the sites were extracted from StreamCat ^74^, while catchment area and slope were extracted via NHDPlusV2.1 with the R package ‘nhdplusTools’^75^. The custom R script was also used to extract aridity data from the Global Aridity Index and Potential Evapotranspiration (ET0) Database: Version 3^76^ with the terra::rast() and terra::extract() functions^77^.

### 2. Sensor deployment and data collection

During the week of July 25–28, 2022, dissolved O_2_ (DO) sensors (miniDOT Logger; PME, Inc.; Vista, CA, USA) and level loggers (HOBO^®^ Water Level Logger (13’) U20L-04; Onset; Bourne, MA, USA) were deployed with sensor windows facing upstream on the thalweg of the riverbed at each site (n = 48) for 35 days during baseflow conditions. We were unable to deploy sensors and sample first order streams due to the lack or low flow during the summer. The sensors were secured to two bricks with zip ties using several different configurations depending on streamflow velocity, depth, and channel substrate (Fig. 7b, d). DO sensors recorded DO and water temperature continuously on a 1-min interval. To minimize sensor biofouling in warmer, shallower rivers or rivers that received a lot of sunlight, 25 DO sensors were equipped with wipers (miniWIPER for miniDOT; PME; Vista, CA, USA) and anti-fouling copper plates (miniDOT Anti-fouling Copper Option; PME; Vista, CA, USA) (Fig 7d). Wipers were programmed to wipe the sensor face once every 12 hours. The remaining 23 DO sensors were equipped with full copper kits, which included a copper ring, silicone ring, and a copper mesh (Fig. 7b). Non-vented level loggers recorded absolute pressure and water temperature continuously on a 15-min interval. Additional barometric pressure (BP, Rugged BaroTROLL Data Logger (9 m); In-Situ; Fort Collins, CO, USA) sensors were deployed at a subset of sites (n = 9) to collect continuous BP and air temperature data throughout the study period (Fig. 7). Sensors were deployed from a 1-m rope on a tree branch as close as possible to the DO sensor/level logger deployments and recorded BP and air temperature continuously on a 1-min interval. Barometric pressure data were used to convert level logger absolute pressure data to depth data.The nine barometric pressure sensors were strategically distributed across different elevations within the basin (340-1,200 m) to capture atmospheric pressure variation. To estimate barometric pressure for sites without a sensor, we fit a regression model between measured barometric pressure and elevation for every 15-min logging interval for all sites with a sensor. In total, we fit ∼3600 linear regression models. These models showed strong correlation between elevation and pressure (R² ranging 0.97-1), providing confidence that we could accurately estimate a pressure time series at sites without sensors using the regression models.

**Figure 7.**
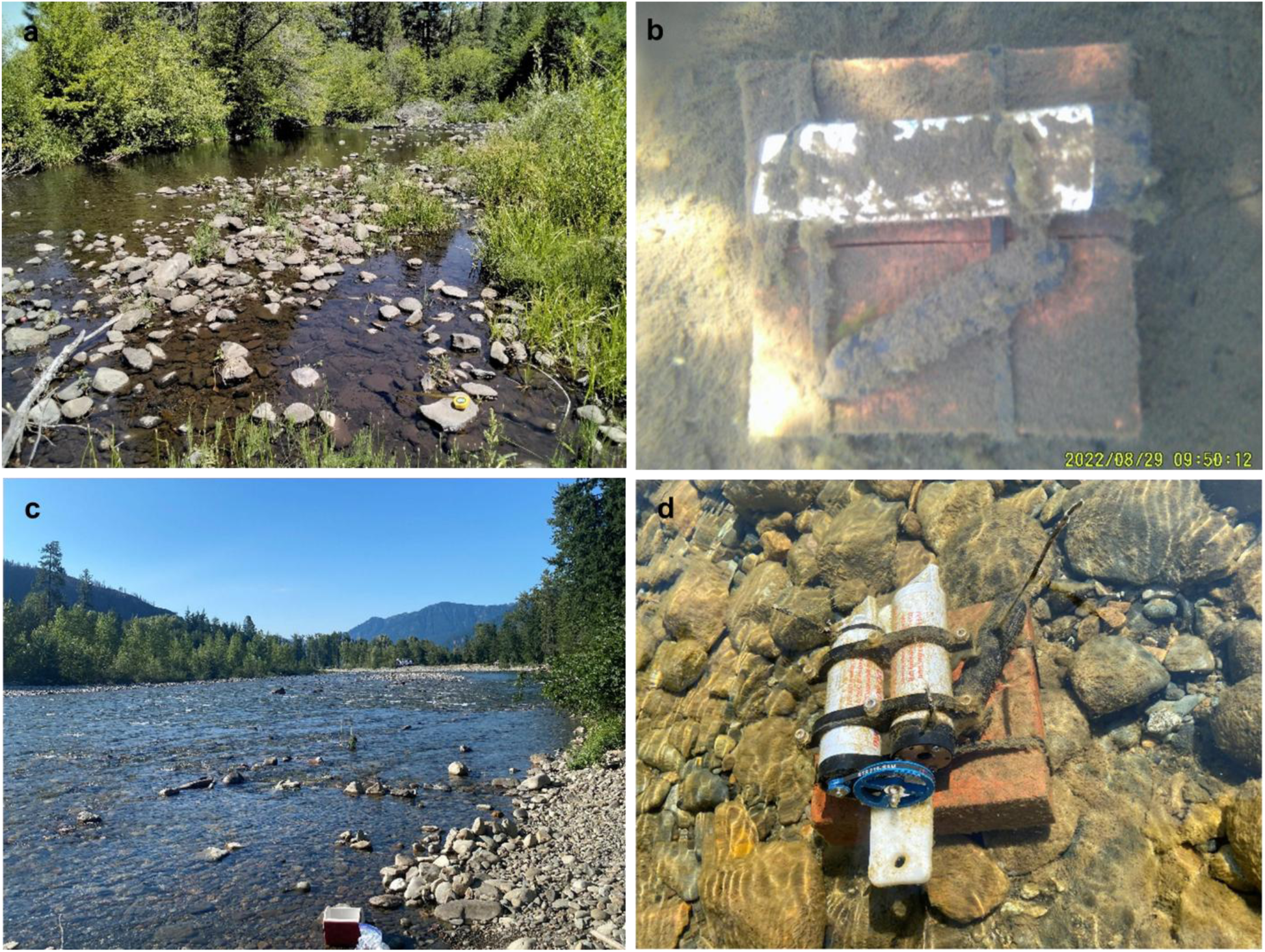
Example sensor deployment configurations on example study sites. Left panels emphasize the variable environmental settings covered in this study. Right panels show two different sensor configurations used to accommodate different environmental conditions. (a) Bull Creek (site S63P), Yakama Reservation, Yakima County, Washington, 25 July 2022. (b) DO sensor and level logger deployed side-by-side on top of two bricks and secured with zip ties, 29 August 2022. DO sensor equipped with an anti-biofouling copper plate and mesh to inhibit the growth of periphyton on the sensor face. Note the dense accumulation of periphyton and sediment deposited on the sensor bodies and bricks. (c) Cle Elum River (site S24R) upstream of Cle Elum Lake, Kittitas County, Washington, 11 August 2022. (d) Level logger (far right) and DO sensor equipped with a wiper kit (white sensor body on left-hand side) and anti-biofouling copper plate, 01 September 2022. The white sensor body on the left-hand side of the wiper kit controls the blue flywheel, which is equipped with a brush and programmed to wipe the sensor face once every 12 h. Note the lighter accumulation of periphyton and sediment on the equipment deployed in the Cle Elum River (d) compared to the equipment deployed in Bull Creek (b).

Level loggers and BP sensors were calibrated by the manufacturer and did not need further calibration. DO sensor calibrations were verified at 100% O_2_ saturation in the laboratory prior to deployment and field-based data were corrected for sensor drift by applying a correction factor using lab-generated data (Supplementary methods). Approximately mid-way through the sampling period (August 8–11, 2022), sensors were field-checked for signs of disturbance, exposure due to decreased flows, and biofouling. DO sensors and level loggers were retrieved temporarily for maintenance and data downloads and immediately redeployed in the original location or, if exposed by decreasing flows, redeployed closer to the channel thalweg. Sensors were retrieved the week of August 29–September 1, 2022, and taken to the laboratory for data download.

### 3. Channel depth, channel width, and discharge data collection

Depth data for sites located on wadeable streams (n = 39) was collected using wading-based depth transects. For each desired reach length (Supplementary methods), we completed 5-10 depth transects over a period of one hour (a minimum of five transects was desired) using 2-person teams, starting ∼2 m upstream of the depth sensor. Transects were evenly distributed along the length of the reach. Using a wading rod, depth measurements were collected along the wetted width of the reach at five roughly equidistance points along a transect perpendicular to flow. Field personnel manually recorded the start time and the coordinates for the starting point of each transect; the end time of the last transect was also manually recorded. The channel width of each transect was measured using a range finder; mean channel width was calculated by taking the average of the range finder width measurements.

Depth data for sites located on non-wadeable rivers (n = 5) was collected using kayak- or jetboat-mounted sonar (HDS-7 LIVE fish finder with Active Imaging™ 3 in 1 transducer; Lowrance; Tulsa, Oklahoma, USA) or, in the case of very large or inaccessible rivers (n = 4), publicly available gauging station data. Kayak-mounted sonar data were collected along each non-wadeable reach by 3-person kayak teams using a sonar receiver mounted to the bottom of a fishing-style kayak. Continuous point measurements were collected by turning on the sonar at the top of each reach and floating downstream using a consistent, zig-zag pattern from bank-to-bank along the entire length of the reach. Coordinates for the top and bottom of each reach and sonar start and end times were manually recorded. HOBO level loggers mounted on a meter stick were used to record depth in reach sections too shallow to collect accurate sonar data.

Mean channel width for non-wadeable rivers where depth data were collected using sonar was calculated for each reach by averaging a minimum of five width estimates from the sonar map. For two sites on the Naches River, WA, river depth and channel width were determined from a rating curve developed by the Washington Department of Ecology (https://apps.ecology.wa.gov/publications/SummaryPages/1803009.html). For three sites on the Yakima River, river depth and channel width were determined from a rating curve developed using data from the United States Geological Survey (USGS) gauging station at Union Gap, Kiona and Mabton.

Stream discharge and velocity for each site were extracted from the National Oceanic and Atmospheric Administration (NOAA) National Water Model Retrospective Dataset Version 2.1 (NWM V2.1; https://registry.opendata.aws/nwm-archive/). This dataset provides a 42-y (February 1979 through December 2020) hourly simulation at all sampling locations. We computed the August monthly average discharge and velocity across these simulation years as input variables for determining an estimate of gas exchange rate (K600) for streamMetabolizer model runs (Section 5). Median grain size (d50) data for all the sites were extracted from NEXSS^11^ which uses d50 data from the National Rivers and Streams Assessment and the Wadeable Stream Assessment (https://www.epa.gov/national-aquatic-resource-surveys/nrsa).

### 4. Surface water sampling

During the week of August 8–11, 2022, surface water samples were collected in triplicate from each site. Surface water samples were collected at mid-water column depth, maintaining consistent relative sampling depth across all sites. A set of samples were collected using a 50-mL syringe and filtered into 40-mL amber glass vials (ThermoScientific™ Amber Clean Snap Vials) using a 0.22-μm sterivex filter (MilliporeSigmaTM Sterivex™ Sterile Pressure-Driven Devices; MilliporeSigmaTM; Burlington, Massachusetts, USA). Filtered samples were analyzed for dissolved organic carbon (DOC) and total dissolved nitrogen (TDN). Additionally, one unfiltered water sample was collected using a pre-washed 2-L amber bottle (Nalgene™ Rectangular Amber HDPE Bottles; ThermoFisher Scientific; Waltham, Massachusetts, USA) for total suspended solids (TSS). All samples were stored on ice in the field and then refrigerated at 4° C before shipping for analysis at the Pacific Northwest National Laboratory (PNNL) Marine and Coastal Research Laboratory in Sequim, Washington (DOC, TDN) or transported to PNNL Biological Sciences Facility Laboratory in Richland, Washington (TSS). TSS samples were analyzed within one week of collection and DOC and TDN content were measured within two weeks of collection.

Dissolved organic carbon (DOC) and TDN were measured using a Shimadzu TOC-L Total Organic Carbon Analyzer. DOC was measured as non-purgeable organic carbon and TDN was measured by converting nitrogen in the samples to nitrogen gas and detected via chemiluminescence. Visual checks of calibration curves, samples, blanks, and check standard peaks were performed before exporting data. Concentrations below the limit of detection (LOD) of the instrument, or below the standard curve, were flagged^78^. The LOD for DOC was 0.17-0.51 mg C L^-1^ and the LOD for TDN was 0.027 mg N L^-1^. If the concentration of an analyte was below the lowest standard but above the LOD, the flag for that sample was replaced with half of the concentration of the lowest standard. If the analyte was below the LOD, the flag for that sample was replaced with a value corresponding to half of the LOD for that run. The coefficient of variation (CV) for each replicate set was calculated. Replicates with CV of 30 % or higher were flagged and the outlier sample was identified by calculating the distance between each of the replicate samples. Replicates with the highest distance to the other two replicates were flagged as outliers and removed from the analysis. Mean values for each site were then calculated from the remaining replicates and input as predictor variables in the regression analysis (Section 8).

Samples for TSS were filtered in the laboratory through a pre-weighed and pre-combusted 4.7 cm, 0.7 µm GF/F glass microfiber filter (Whatman™ glass microfiber filters, Grade 934-AH®; MilliporeSigma; Burlington, Massachusetts, USA). After water filtration, the filter and filtration apparatus were rinsed with 30 mL of ultrapure Milli-Q water (Milli-Q® IQ Water Purification System; MilliporeSigma; Burlington, Massachusetts, USA) to ensure that all residue was captured by the filter. The filter was placed in foil and oven dried overnight at 45° C. The filter was allowed to cool in a desiccator prior to weighing. TSS was calculated as the difference between the weight of the filter before and after filtration of the water sample divided by the volume of water filtered^79^. Similar to the approach applied to DOC/TN, if the TSS of a sample was below the LOD, the flag for that sample was replaced with a value corresponding to half of the LOD (TSS LOD = 0.24 mg L^-1^) for the TSS method.

### 5. Total Ecosystem Respiration (ER_tot_) estimates

We estimated ER_tot_ using the streamMetabolizer R package^25^ which estimates ER, GPP and gas exchange (K600) as parameters fit to a diel model of DO. Model inputs consisted of time series data for DO, water temperature, BP, adjusted time series of depth (Supplementary methods), site ID, site latitude and longitude and a site-specific prior estimate of gas exchange rate coefficient (K600, day⁻^1^). Gas exchange normalized to a Schmidt number of 600 (K600, day⁻^1^) is a free parameter estimated within streamMetabolizer via Bayesian inference and therefore requires a prior probability estimate. We first estimated a prior k600 (m day⁻^1^) value using the empirical relationship described in Model Equation 7 from Table 2 in Raymond et al. (2012)^80^ as well as site specific slope, discharge, and velocity (Section 1 and 3). k600 values were then divided by the reach-averaged depth (Supplementary methods) to obtain a prior probability estimate for K600.

Before running the model, the 15-min time series BP and reach-averaged depth were interpolated (zoo::na.approx function)^81^ to match the time interval of the water temperature and DO. Then the temperature, DO, and BP 1-min time series data were down-sampled to a 15-min resolution. Down-sampling improved both solution time and K600 pooling. DO time series was visually inspected and days with aberrant DO signals or biofouling were removed from the timeseries prior running the model. Additionally, we flagged and interpolated outliers in the DO time series using the R function ‘auto_data_cleaning’ within the ‘tsrobprep’ package^82^ and used interquartile range for outlier detection. Further, photosynthetically active radiation (PAR) was estimated using the streamMetabolizer::calc_light() function^25^, which is based on solar time, latitude, and longitude. Saturated DO concentration was calculated for each DO time series using water temperature and BP times series data and the Garcia and Gordon (1992)^83^ equation (streamMetabolizer::calc_DO_sat function)^25^.

For most of the sites, the Bayesian model was configured with normal K600 pooling, which is to say we partially pooled daily log K600 estimates in a Bayesian multilevel model. We also used the state-space option in streamMetabolizer to estimate observation and process error for DO. The prior for K600 (K600_daily_meanlog_meanlog) was set as the natural log of the K600 prior estimate and a K600 scale value of 0.7. Group level variation in log K600 was K600∼normal (0,0.05). We estimated posterior probability distributions for parameters via Hamiltonian Monte Carlo using the program Stan with 2000 warm-up steps, and 2000 saved steps across four chains. We assessed model convergence by evaluating Rhat and n_eff distributions. A model demonstrated good convergence when n_eff > 100 and Rhat ≤ 1.1. The K600 daily sigma prior was adjusted (0.08, 0.02, 0.01, or 0.05) on a site-by-site basis to optimize model performance. StreamMetabolizer outputs comprised ER_tot_ (g O_2_ m⁻^2^ day⁻^1^), GPP (g O_2_ m⁻^2^ day⁻^1^), and K600 (day⁻^1^) as mean daily estimates over the number of days with quality DO data over deployment time^84^. Finally, we calculated the mean GPP and ER_tot_ for the entire deployment time for each site.

### 6. Water column respiration (ER_wc_) and sediment-associated respiration (ER_sed_) estimates

We estimated water column respiration in triplicate for 75 min at each site during the sampling week using the *in situ* dark bottle incubation approach previously described by Laan et al.^33^.

Briefly, a DO sensor (miniDOT Logger; Precision Measurement Engineering, Inc., Vista, CA, USA) and a small, battery-powered mixing device (i.e., toy boat motor propellor) were placed inside a 2-L dark bottle in triplicate (Nalgene™ Rectangular Amber HDPE bottles; ThermoFisher Scientific, Waltham, Massachusetts, USA). DO sensors and mixing devices were placed in a cooler with blue ice packs to keep them cool and minimize the time needed at each site for the sensors to equilibrate with river water temperatures. Bottles with the sensor and mixing device were rinsed three times with river water, and filled with river water from the thalweg, ensuring no air bubbles were trapped inside. Once full, the bottles were deployed on the riverbed as close to the thalweg as possible, depending on river depth and access to the shoreline, and weighted down with bricks or larger rocks. The sensors recorded DO concentration and temperature at 1-min intervals during the 75-min incubation period.

We extracted the raw DO and temperature sensor data for each site and plotted each variable against incubation time for each set of triplicate incubations (n = 144). The plots were visually inspected to a) confirm that temperature sensors were at equilibrium with the river temperature when the 75-min incubation test period began and b) identify data gaps, outliers, and other data anomalies. Following the visual inspection of plots, we applied the DO correction factor (as described in Supplementary Methods), removed the first 5 minutes and trimmed the time-series to 70 min to account for data anomalies due to emptying and refreshing river water in the bottles at the beginning and end of the incubation period, and to ensure all sites had the same incubation time.

Water column respiration rates (ER_wc_) for individual triplicate incubation samples were calculated as the slope of the linear regression model (R stats::lm function)^85^ between DO and incubation time in units of day⁻^1^. The volumetric daily rate was then converted to daily areal units (g O_2_ m⁻^2^ day⁻^1^) by multiplying by the reach-averaged depth (Supplementary methods). Only sample observations with a normalized root mean square error (NRMSE) ≤ 0.03 were used for further analysis^86^. Although all triplicate samples met the NRMSE criteria, we removed a replicate from two sites after visual inspection due to the curve deviating from the other two replicates. Furthermore, six additional sites only had two replicates due to sensor or data loss. Median, mean, and range of ER_wc_ values for each site were calculated. 40% of ER_wc_ values were slightly positive (i.e., ER_wc_ > 0). Although positive respiration rates are biologically unrealistic, and values greater than 0 g O_2_ m⁻^2^ day⁻^1^ but less than 0.5 g O_2_ m⁻^2^ day⁻^1^ are difficult to distinguish from zero, we retained positive ER_wc_ values to prevent biasing the data.

Respiration directly and indirectly associated with sediments (ER_sed_) was found by the difference between ER_tot_ and ER_wc_, assuming zero ER_wc_ if the estimated values were positive. These ER_sed_ estimates include sediment-associated respiration from all ecosystem components besides the water column.

### 7. Model-predicted hyporheic zone respiration

Model-predicted HZ respiration values were previously published in Son et al., (2022)^14^, and the data are publicly available on ESS-DIVE^87^. Respiration values by Son et al. (2022)^14^ correspond to the amount of aerobic respiration in the HZ via vertical and lateral hyporheic exchange between the stream and the HZ. The respiration is scaled by the stream surface area.

Respiration values reported in Son et al., (2023)^87^ in units of mol O_2_ day⁻^1^, were converted to g O_2_ m⁻^2^ day⁻^1^ using the molar mass of O_2_ and RCM stream reach area. We note that model outputs cover only 46 out of the 48 sites that we studied in the YRB.

### 8. Data analysis

To explore how physical variables covary with ER_sed_ at different sites, a LASSO (Least Absolute Shrinkage and Selection Operator) regression model was built using physical, chemical and environmental variables (Table 4) as inputs, and ER_sed_ as the response variable. We used drainage area as a quantitative variable in the LASSO analysis, as opposed to categorical stream order, because it is a continuous variable representing stream characteristics across the basin. Prior to conducting the LASSO regression, variables were cube root transformed to reduce the impact of high leverage points in the regression analysis. Additionally, standardization was applied to achieve a mean of zero and standard deviation of 1 (i.e., z-score) to equally weight all variables. The LASSO regression was performed over 100 iterations, each with a different random seed. β coefficients were normalized to the maximum β coefficient for each iteration, then averaged over the 100 iterations for the final reported value. Both the raw and normalized mean β coefficient and standard deviation are reported in addition to the R^2^ (Table 2).

**Table 4.** Variables used as inputs in the LASSO regression model. All NHDPlus, StreamCat, NLCD, and ET0 database data were extracted using a custom R script^73^.

| Variable | Description | Unit | Data Source | Data Source Citation |
| --- | --- | --- | --- | --- |
| Water Temperature | HOBO water temperature | C° | Field measured | Delgado et al. (2023)<sup>79</sup> |
| Mean Depth | average water depth | cm | Field estimated | Delgado et al. (2023)<sup>79</sup> |
| Slope | stream slope | NA | NHDPlus | McKay et al. (2012)<sup>75</sup> |
| Velocity | stream velocity | m s<sup>-1</sup> | National Water Model | Cosgrove et al. (2024)<sup>88</sup> |
| Discharge | Discharge | m<sup>3</sup> s<sup>-1</sup> | National Water Model | Cosgrove et al. (2024)<sup>88</sup> |
| Dissolved Organic Carbon | non-purgeable organic carbon | mg L<sup>-1</sup> | Field measured | Forbes et al. (2023)<sup>78</sup> |
| Total Dissolved Nitrogen | total dissolved nitrogen | mg L<sup>-1</sup> | Field measured | Forbes et al. (2023)<sup>78</sup> |
| Total Suspended Solids | total suspended solids | mg L<sup>-1</sup> | Field measured | Delgado et al. (2023)<sup>79</sup> |
| Drainage Area | total drainage area | km<sup>2</sup> | NHDPlus | McKay et al. (2012)<sup>75</sup> |
| Percent Forest Cover | percentage of forest as the sum of percentage deciduous, coniferous and mixed forest | % | StreamCat and NLCD | Hill et al. (2016)<sup>74</sup><br>Jin et al. (2023)<sup>89</sup> |
| Percent Agricultural Cover | percentage agriculture as the sum of percentage crop and hay | % | StreamCat and NLCD | Hill et al. (2016)<sup>74</sup><br>Jin et al. (2023)<sup>89</sup> |
| Percent Shrub Cover | Percentage Shrub | % | StreamCat and NLCD | Hill et al. (2016)<sup>74</sup><br>Jin et al. (2023)<sup>89</sup> |
| Aridity | Aridity | NA | ET0 Database | Zomer et al. (2012)<sup>90</sup> |
| Median grain size (d50) | median grain size | m | National Rivers and Streams Assessment and the Wadeable Stream Assessment | Kaufman et al. (2023)<sup>84</sup> |
| Gross Primary Production (GPP) | gross primary production | gO<sub>2</sub> m<sup>-2</sup>·day<sup>-1</sup> | streamMetabolizer output | Kaufman et al. (2023)<sup>84</sup> |

We used LASSO regression for exploratory analysis to identify which variables collectively explain variation in ER_sed_ rather than to test specific hypotheses about individual variable effects. LASSO automatically performs variable selection by shrinking unimportant coefficients to zero while preventing overfitting that commonly occurs with traditional stepwise selection for cases with co-correlated independent variables. This is useful because we had 15 explanatory variables, some being strongly co-correlated. LASSO also emphasizes relatively simple models and validates results through cross-validation rather than p-values, assessing model performance across multiple data subsets to identify variables that consistently contribute to predictive accuracy.

To investigate the correlation between z-score normalized RCM-predicted HZ respiration and z-score normalized field-estimated ER_tot_ and ER_sed_, Pearson and Spearman rank-order correlations were used. Rank-order correlation was used when data violated normality assumptions of parametric analysis. In addition, the linear regression analyses between GPP and ER_tot_/ER_sed_ (Fig. 5) were performed using the ‘lm’ function in the base R package. All analyses were performed using R Statistical Software (v4.1.3)^85^.

## Data Availability

Data collected in the field were published previously in Forbes et al. (2023)^78^ and Delgado et al. (2023)^79^. Data generated as part of this manuscript are currently available on GitHub (see below) and will be published on the Environmental System Science Data Infrastructure for a Virtual Ecosystem (ESS-DIVE) repository upon manuscript acceptance. ESS-DIVE repositories are licensed for reuse under the Creative Commons Attribution 4.0 International License.

## Code Availability

Scripts necessary to reproduce the primary results of this article are available at https://github.com/river-corridors-sfa/SSS_metabolism.

## Acknowledgements

This work was supported by the River Corridor Science Focus Area (RCSFA) at the Pacific Northwest National Laboratory (PNNL). The RCSFA is supported by the United States Department of Energy, Office of Biological and Environmental Research (BER), Environmental System Science (ESS) Program. PNNL is operated by Battelle Memorial Institute for the United States Department of Energy under contract no. DE-AC05-76RL01830. We thank the United States Forest Service, Washington Department of Natural Resources, Washington Department of Fish and Wildlife, Confederated Tribes and Bands of the Yakama Nation, and Cowiche Canyon Conservancy for access to field locations where these samples were collected. We also thank the Confederated Tribes and Bands of the Yakama Nation Tribal Council and Yakama Nation Fisheries for working with us to facilitate sample collection and optimization of data usage according to their values and worldview. We thank the field team including: Dillman Delgado, Morgan Barnes, Brandon T. Boehnke, Yunxiang Chen, Kali Cornwell, Brianna I. Gonzalez, Samantha Grieger, Glenn E. Hammond, Peishi Jiang, Bing Li, Zhi Li, Xinming Lin, Sophia A. McKever, Maruti K. Mudunuru, Katherine A. Muller, Opal Otenburg, Aaron Pelly, Kelsey Peta, Alan Roebuck, Joshua M. Torgeson, and Jianqiu Zheng.

## Author Contributions

VAGC, MK, BH and JCS conceptualized the study. JCS, LR, SF, ML conducted the field work. LR and ML analyzed samples. KS, XC and YF configured RCM outputs for this study. XL, ML and BF processed the data and performed analyses with feedback from VAGC, MK, BH and JCS. VAGC and MK carried out metabolism analysis with feedback from BH. BF managed the data. VAGC, MK, SF, BF and JCS drafted the initial manuscript, and all authors contributed to revisions.

