## Supplementary Information for "Sediment-associated processes account for most of the spatial variation in stream ecosystem respiration in the Yakima River basin"

**Supplementary methods**

1. Data processing

Data processing and analyses were completed in R[1]. Most data cleaning and manipulation prior to analysis was done using the tidyverse[2] R package. .

a. Dissolved oxygen sensor time series

Sensor calibrations were verified in the laboratory following manufacturer recommendations via a bubbler test. The bubbler test was conducted in groups of 10 sensors by immersing them in a 5-gal bucket of tap water (sensor window facing up with a minimum of 15–20 cm of water covering the window) 100% saturated with ambient air using an aquarium pump and air stone. Continuous DO concentration, water temperature and local barometric pressure data were logged at 1-min intervals for 24 h. Using a gas calibration system[3], we found the water in to bucket to be over saturated by 1% and added this correction factor to our processing steps. In turn, we calculated maximum theoretical DO concentration ($DO_{sat-max}$) in saturation conditions at the laboratory pressure and water temperature via the Garcia and Gordon[4] equations at each time step. $DO_{sat-max}$ was multiplied by 0.99 to account for the 1% oversaturation. The sensor specific correction factor was then calculated as the average of the ratio between measured DO concentration and $DO_{sat-max}$ over the 24 h of data collection. We applied the sensor specific correction factor to the data from each sensor to account for sensor drift. Nineteen out of 63 DO sensors fell outside of the manufacturer's calibration recommendations (± 5% of $DO_{sat-max}$; i.e., $0.95 < DO_{sat-max} < 1.05$).

b. Reach-averaged depth time series

We calculated reach-averaged depth to account for the influence of the entire reach length on the DO signal at the deployment location. We defined a reach length as approximately 50 times the width of the stream. We then estimated depths via wading-based depth transects, sonar, or publicly available depth along this approximate reach length. Further, a single value for reach-averaged depth was calculated at each site by summing all observed depth measurements along the reach and dividing them by the total number of observations. This value corresponded to a reference measured reach-depth at single-point-in-time which was then used to correct the time series measurements of depth, which were collected at one point location for each reach throughout the sensor deployment.

Reach-averaged depth time series were created for each site using pressure and temperature from a HOBO level logger along with barometric pressure, air temperature and reach-averaged depth at a single-point-in-time from each site. As was done with the DO sensor time series datasets, we merged the two pre- and post-sampling week raw level logger time series datasets into a single, continuous time series dataset for each site; plotted each parameter over time; visually inspected the plots for data gaps, outliers, and other data anomalies; and removed spurious data recorded immediately before and after sensor deployments and following sensor retrieval at the end of the monitoring period. To account for changes in the deployment location of the sensor following sampling week redeployments, we calculated an offset subtracting the pressure before removal from pressure after redeployment and then subtracted this offset value from the timeseries after redeployment. In addition to post-sampling-week data changes, 8 sites had jumps in data (rapid decrease or increase) that could have been due to the sensor moving as a result of water movement or from human interference. We determined that these jumps were not caused by weather or dam releases, so we corrected the data calculating and applying an offset in the same way as it was done after redeployment. We then atmospherically corrected HOBO pressure data by subtracting local barometric pressure at each time-step. Furthermore, no data interpolation was needed to fill time gaps in the time series due to both a) the short length of time that the sensors were out of the water and b) the 15-min time increments.

Following pressure corrections, a time series of depth at each site was estimated via the hydrostatic equation as the quotient between the corrected pressure and the density of water times gravity. Density of water was calculated at site conditions using Equation 6 in Jones and Harris[5] and the time series of water temperature. Further, we calculated a depth offset as the difference between the reach-averaged depth at single-point-in-time and the sensor estimated depth at the time the reach-average depth was collected. Finally, the reach-averaged depth time series was calculated after subtracting the offset value from each point of the sensor estimated depth time series. For any period with missing data, we used the last observation carried forward (LOCF) interpolation approach (tidyr::fill function)[7] to fill in the missing values. The obtained dissolved oxygen, barometric pressure, and depth time series with 15-minute resolution are used as an input parameter in streamMetabolizer[8] (Section 5 in Methods).

### Supplementary figures

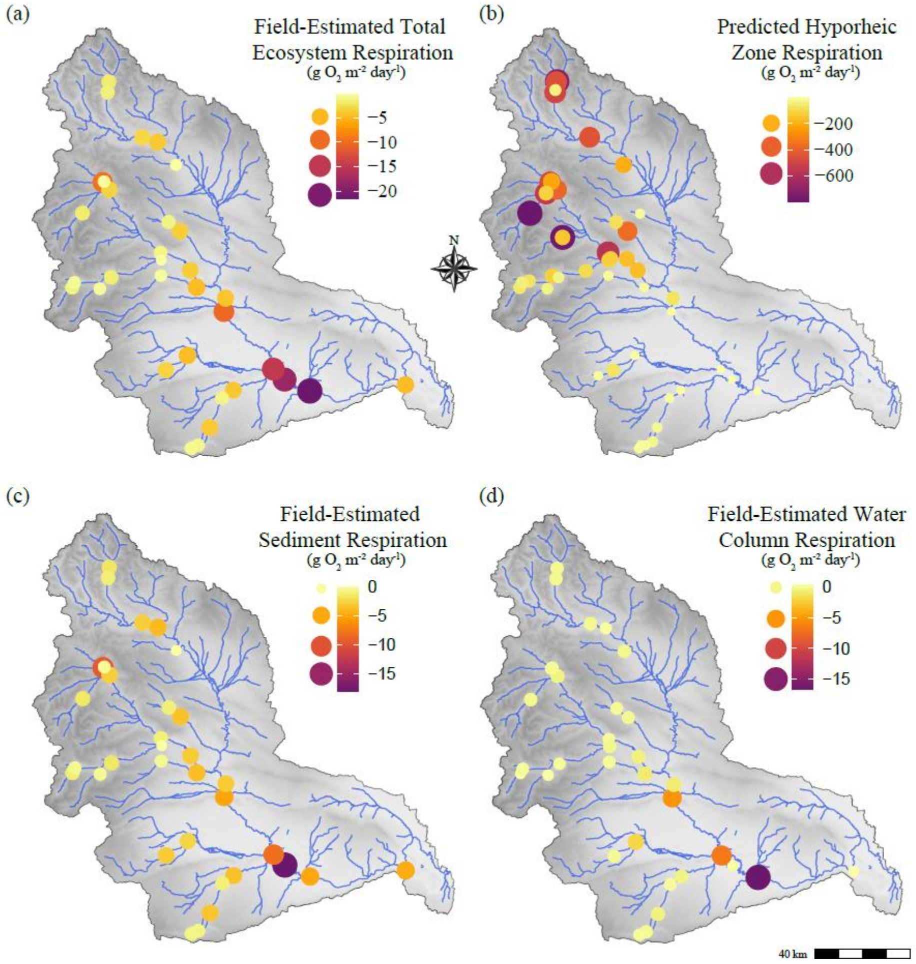


**Figure S1. Map displaying Yakima River basin sites investigated in this study and associated respiration rates. This figure displays the information from Figure 1 in the main text without points where $ER_{sed}/ER_{tot}$ was positive or could not be calculated.** (a) Field-estimates (n = 33) of total stream ecosystem respiration rates ($ER_{tot}$). (b) River Corridor Model (RCM, n = 46) predicted hyporheic zone (HZ) respiration rates[13]. (c) Field-estimates (n = 33 of sediment-associated respiration rates ($ER_{sed}$). (d) Field-estimates (n = 48) of water column respiration rates ($ER_{wc}$). Color and symbol size are used to indicate magnitude of respiration rate.

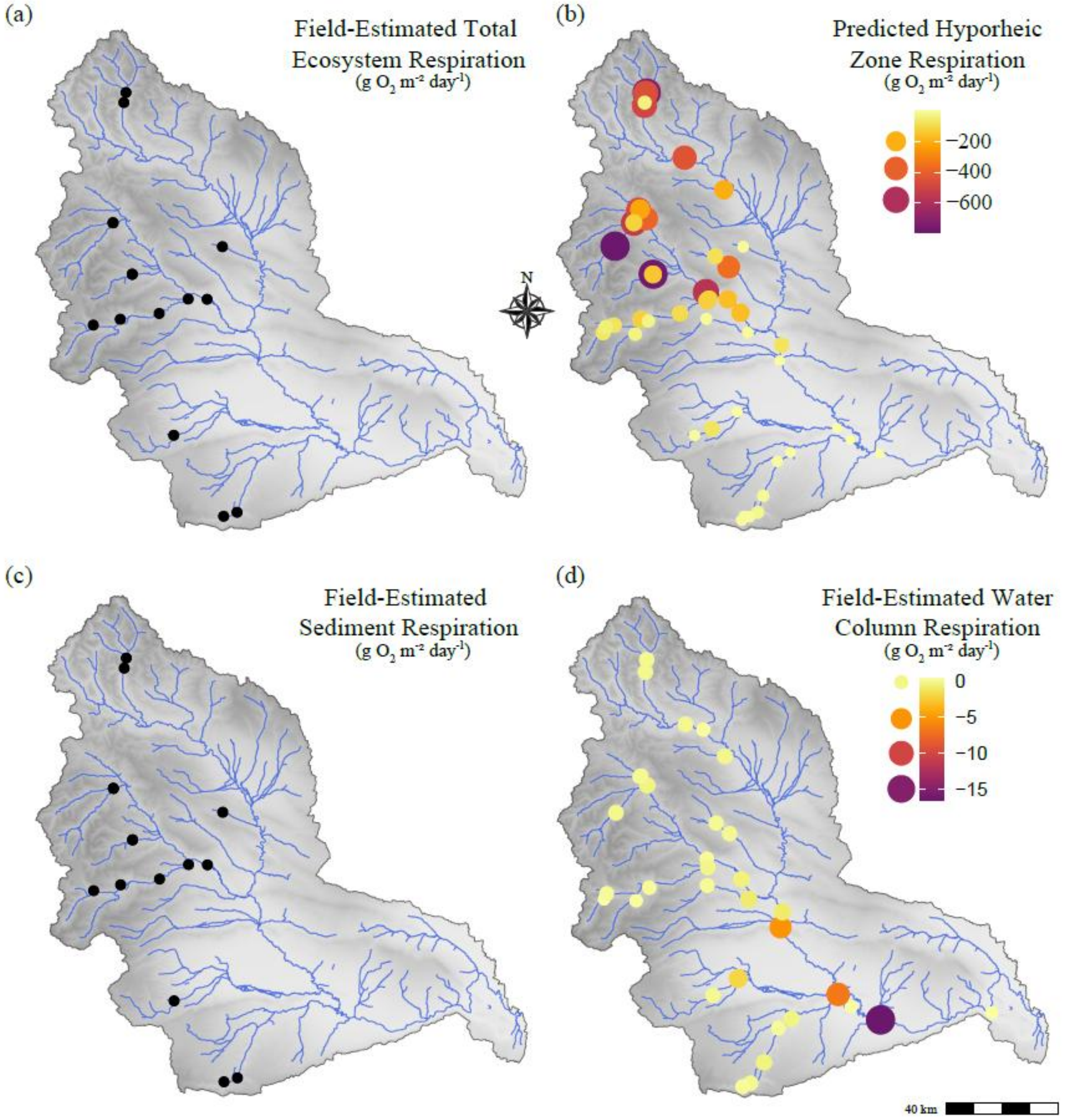


**Figure S2. Map displaying Yakima River basin sites investigated in this study and associated respiration rates.** (a, c) Sampling points where $ER_{sed}$ or $ER_{tot}$ were positive or could not be calculated (n = 15). (b) River Corridor Model (RCM, n = 46) predicted hyporheic zone (HZ) respiration rates[13]. (d) Field-estimates (n = 48) of water column respiration rates ($ER_{wc}$). Color and symbol size are used to indicate magnitude of respiration rate.

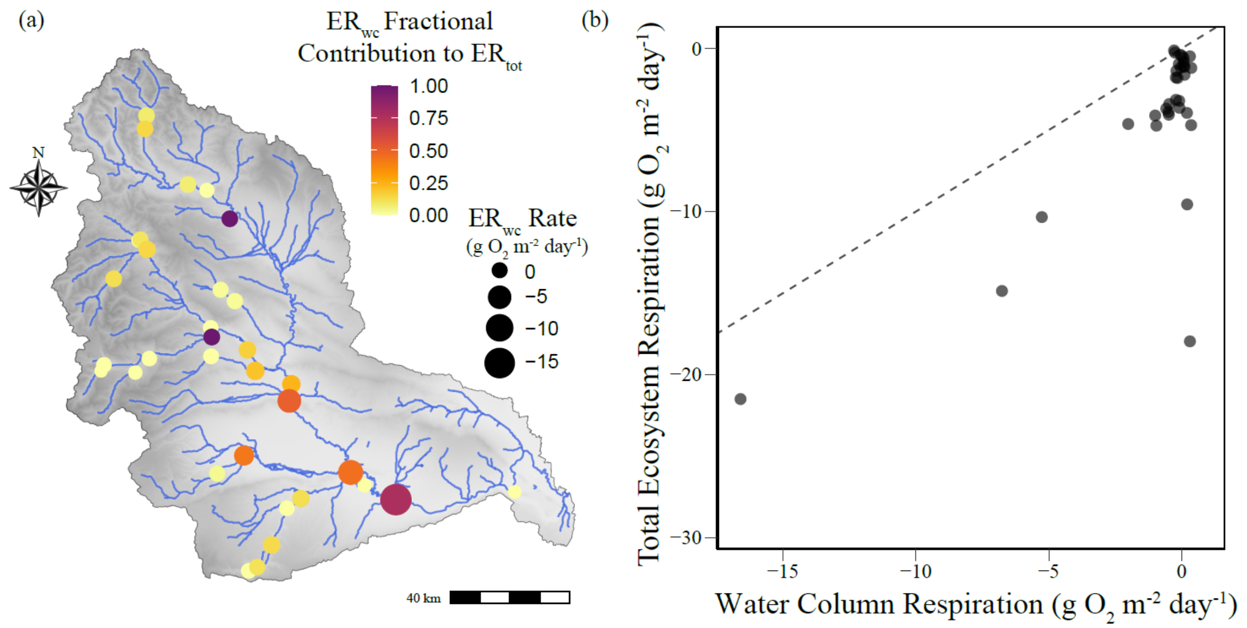


**Figure S3. ER$_{wc}$ contributions and correspondence with ER$_{tot}$.** (a) Map displaying the fractional contributions of water column respiration (ER$_{wc}$) to total ecosystem respiration (ER$_{tot}$) across the YRB (n = 33). Color is used to indicate the magnitude of the fractional contribution while symbol size is used to indicate magnitude of ER$_{wc}$ respiration rate. (b) Scatterplot between ER$_{wc}$ (x-axis) and field estimates of ER$_{tot}$ (y-axis). The grey dashed line is the one-to-one line with slope of –1.

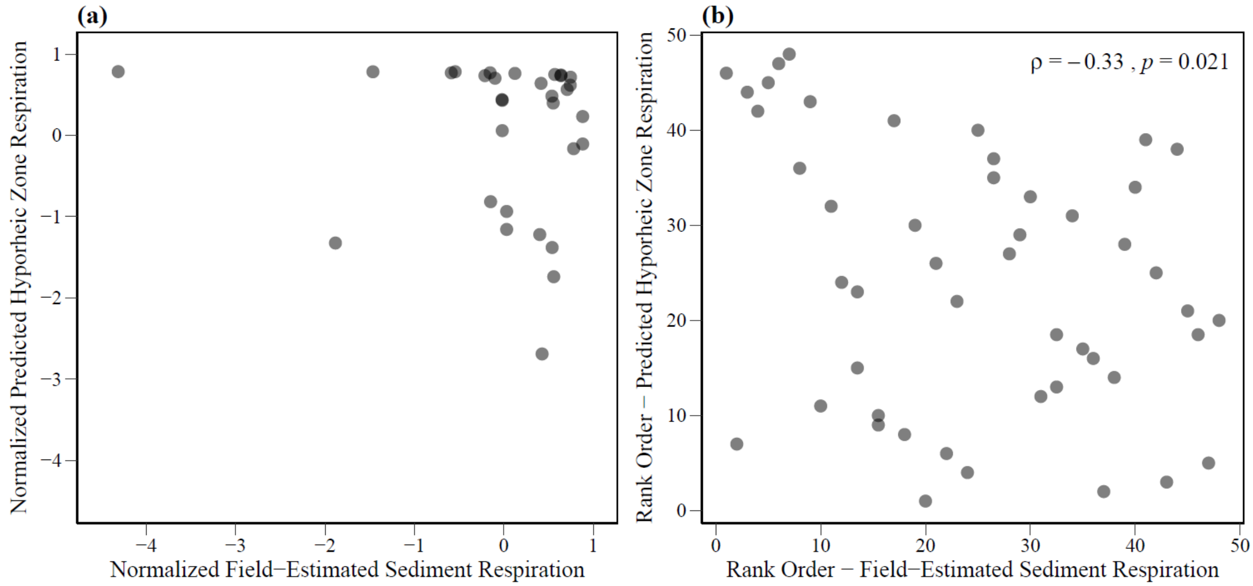


**Figure S4. Normalized model-predicted hyporheic zone (HZ) respiration rates do not increase with field-estimates of sediment associated respiration (ER$_{sed}$; n = 31).** (a) Z-score normalized data suggest no relationship. Data distributions are skewed (i.e., not normal) and thus violate assumptions of parametric regression, so no statistics are given. (b) Spearman rank-order correlation reveals a weak, but significant, negative relationship. The Spearman correlation statistics are provided on the panel.
